# Altered Use of Prior Expectations and Modified Neural Dynamics in a Mouse Model of Autism

**DOI:** 10.1101/2025.09.12.675910

**Authors:** M. Felicia Davatolhagh, João Couto, Max Melin, Lukas T. Oesch, Charles Findling, Polo Morales, Isabelle Hong, International Brain Laboratory, Anne K. Churchland

## Abstract

In dynamic environments, updating beliefs based on past experiences (priors) is essential for optimal decision-making. Prior utilization is often impaired in psychiatric disorders, affecting perception and behavior. We investigate how Neurexin1α (Nrxn1 α) loss-of-function disrupts this process, providing insight into circuit deficits underlying sensorimotor dysfunction. While the synaptic role of Nrxn1α role is well studied, its impact on network dynamics and decision-making behavior remain unclear. Using widefield calcium imaging, we assess cortex-wide activity in mice performing a two-choice task to probe how priors influence visually-guided decisions. This task requires the mouse to combine sensory evidence with the prior probability over the stimulus side. We find Nrxn1α KO mice underutilized priors and were slower to update choices based on feedback. During decision-making, cortex-wide cortical activity is both elevated and increasingly correlated in Nrxn1α KO mice, independent of task period. Moreover, a larger fraction of cortical variance was explained by movement variables, consistent with stronger coupling of cortical activity to motor signals and a bias toward movement-related dynamics. These findings suggest that core computations underlying decision-making, such as integrating past experience with current evidence, depend on intact synaptic mechanisms shaped by genes like Nrxn1α.

## Introduction

Atypical sensory processing is a prevalent feature of neurodevelopmental and psychiatric disorders. Sensory dysfunction frequently emerges as over-or under-reactivity to stimuli, impacting daily functioning ^1^. However, it has been recognized that sensory deficits extend beyond processing, affecting perceptual decision-making as well. Notably, impairments in learning and updating priors (existing beliefs about the probability that certain events will occur), have been observed in humans diagnosed with autism spectrum disorder (ASD) ^2-5^ and in schizophrenia ^6,7^. In optimal decisions, priors are combined with sensory evidence: for instance, a cloudy day (evidence) and the knowledge that one is currently in rainy Portland (prior) might lead to the decision to bring an umbrella to work. These priors are learned: a different decision might be made in response to cloudy weather in California, where the baseline probability of rain is much lower. Given that evidence and priors are combined to evaluate sensory stimuli, and are especially critical for uncertain stimuli, an impairment in the use of priors can have profound implications on how sensory information is perceived and integrated to guide decision-making. While significant progress has been made in mapping sensory-related brain circuits, the link between genetic mutations and the neural network changes underlying deficits in perceptual decision-making remains unclear.

To understand how genetic disruptions give rise to altered perceptual computations, it will be important to bridge the gap between molecular-level perturbations and the large-scale cortical dynamics that support prior-based decision-making. Mice with mutations in Neurexin1α (Nrxn1α), a synaptic cell adhesion molecule with broad neuropsychiatric disease association, serve as an entry point for investigating how specific molecular disruptions give rise to systems-level dysfunction. Previous studies have shown Nrxn1α loss-of-function to produce distinct, context-dependent synaptic phenotypes, and behavioral studies have reported changes in goal-directed behavior in mutant mice^8^. Existing work focusing on other genetic models has begun to uncover differences in the ability of ASD genetic models to update their decision-making strategy as conditions change^9^. However, this work has left open questions about how cortical regions interact dynamically to support prior-based perceptual decision-making, and whether disruptions to these interactions drive the altered decision-making in genetic models of ASD.

In this study, we aim to investigate how genetic disruption of synaptic signaling alters the integration of prior expectations and sensory evidence during decision-making. To do this, we trained Nrxn1α KO mice and their wild-type (WT) littermates on a two-choice visuomotor decision-making task designed to leverage sensory evidence with learned prior probabilities. This task allows us to quantify how subjects adapt their choices in response to shifts in stimulus probability, providing a behavioral readout of prior utilization. We found that Nrxn1α KO mice show impaired use of prior information, instead relying more heavily on sensory inputs - resulting in less flexible, feedback-insensitive behavior. Using widefield calcium imaging across the dorsal cortex, we simultaneously captured cortex-wide neural dynamics as animals engaged in the task. This approach enabled us to investigate how expectations are represented in neural population activity, and how these representations relate to behavioral choices. Together, our findings link a well-characterized molecular perturbation to altered decision making behavior and distributed cortical dynamics, offering insight into how disrupted synaptic signaling may contribute to the cognitive features observed in neurodevelopmental disorders.

## Results

### Nrxn1α mice reliably leverage sensory evidence to guide decision-making

To examine the impact of the loss of Nrxn1α during decision-making, we trained both wild-type (WT) and knockout (KO) mice on a head-fixed visuomotor task in which a grating stimulus appeared on the left or right side of a screen, and animals reported its location by turning a wheel (Figure 1A, 1B) ^10^. The animals are first trained on an unbiased version of the task (50:50 stimulus side probability) progressing through multiple stages of training where they are gradually introduced to difficult contrasts.

**Figure 1.**
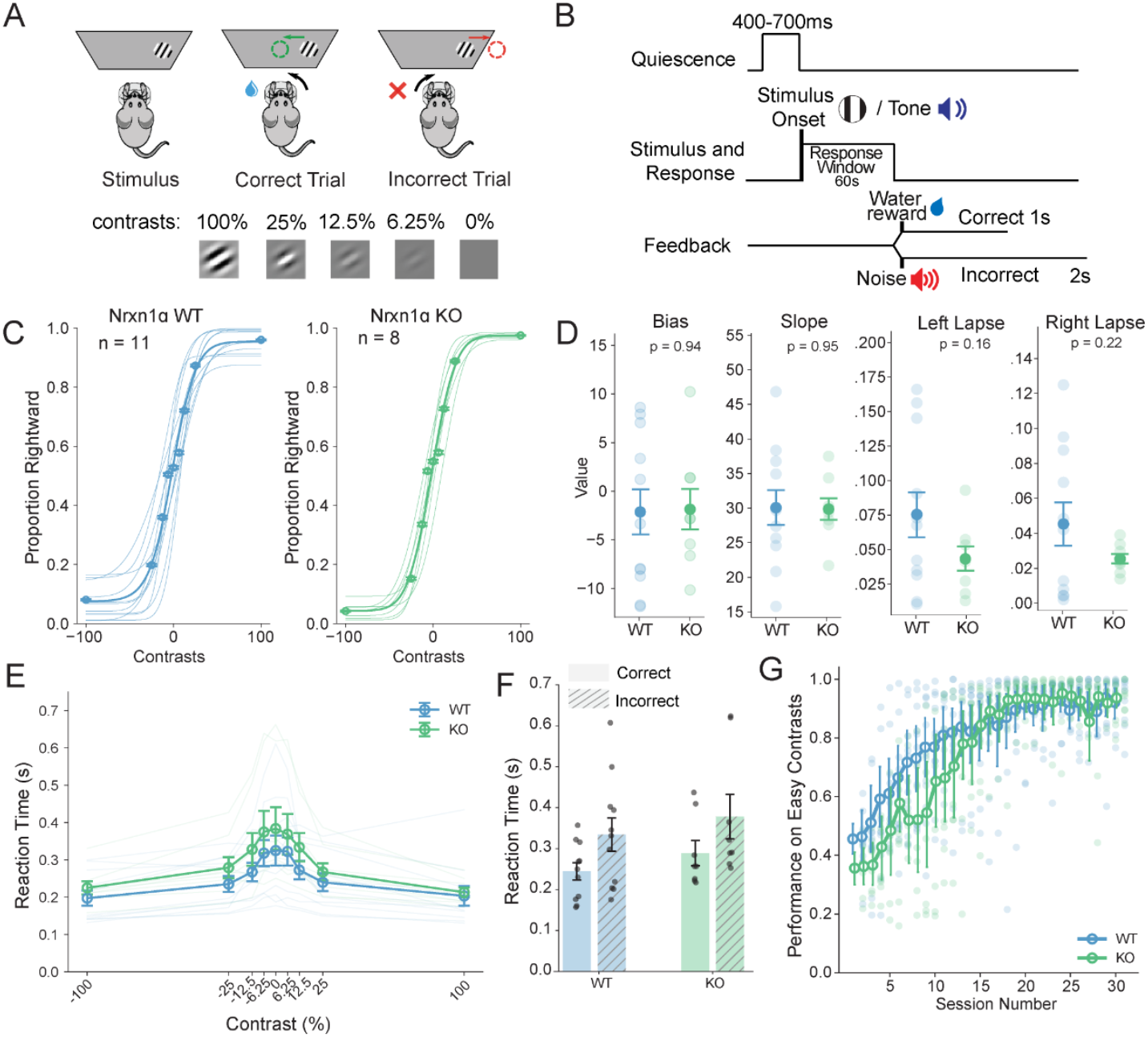
Nrxn1α mice reliably leverage sensory evidence to guide decision-making. **(A)** Schematic of behavioral task depicting stimulus onset, correct trial where stimulus is centered, and incorrect trial where stimulus is moved off-screen. Stimulus contrasts varied across trials. **(B)** Schematic of the trial structure during the visual decision-making task. Each trial began with a quiescence period, followed by stimulus onset. Mice responded by turning a wheel, triggering auditory feedback: a correct response resulted in a reward and tone, while an incorrect response triggered a noise and timeout. **(C)**Psychometric curves showing the proportion of rightward choices across contrast levels for Nrxn1α WT (left, blue) and KO (right, green) mice. Thin lines show individual animals; bold lines show group means with SEM. 10 sessions per animal, WT: 11 mice, KO: 8 mice. **(D)** Summary of psychometric parameters including bias, slope, and lapse rates. No significant differences were observed between WT and KO mice. **(E)** Time to first movement (reaction time) as a function of contrast. Both genotypes showed longer latencies at lower contrasts. **(F)** Average reaction times for correct and incorrect trials, grouped by genotype. **(G)** Learning curve showing performance on easy contrast trials (≥50%) across training sessions. Both WT and KO mice improved over time, with similar overall trajectories.

We first asked whether KO animals were capable of learning the task and achieving asymptotic performance. To evaluate decision-making performance in WT and KO animals, we compared their psychometric functions, which allow dissociation of changes in sensitivity from those in bias or lapse rate ^11^. Both WT (n = 11) and KO (n = 8) animals exhibited typical sigmoidal psychometric curves (Figure 1C), with an increasing likelihood of rightward choices as stimulus contrast increased in that direction. Quantification of psychometric parameters (Figure 1D) showed no significant differences between WT and KO mice in bias (*p* = 0.94), slope (*p* = 0.95), left lapse (*p* = 0.16), or right lapse (*p* = 0.22). We next examined reaction time, finding no differences between genotypes (Figure 1E) and between correct (*p* = 0.234) and incorrect (*p* = 0.522) choices (Figure 1F). Finally, we asked whether Nrxn1α deletion affects the rate at which the animals learn the task. Analysis of performance on high-contrast trials across training days revealed that both WT and KO mice improved over time with similar learning trajectories, reaching asymptotic performance after ∼20 sessions (Figure 1G, Supplementary Figure 1). Together, these results demonstrate that Nrxn1α KO mice can learn the visuomotor decision-making task, use visual evidence to guide decisions, and initiate appropriate motor responses, thereby establishing a behavioral foundation for subsequent investigation of more subtle deficits in flexible behavior and cortical processing.

### Nrxn1α mice exhibit slower adaptation to shifting reward contingencies

Once animals have established basic task proficiency, they progress to a biased version of the task where animals encounter blocks alternating with stimulus probabilities of 80:20 (left stimulus likely) or 20:80 (right stimulus likely), requiring them to adapt their responses based on these probabilistic cues (Figure 2A). When the contrast of the stimulus is absent or low, the sensory signal is uncertain, so animals should rely on their prior knowledge of the block structure (i.e. the expected side based on recent trials) rather than sensory information alone. This reliance on priors is demonstrated in WT animals by a pronounced shift in the change of rightward choices conditioned by block-type (right-likely - left-likely blocks) and in their psychometric curves towards the higher probability stimulus side (Figure 2B, C). However, Nrxn1α KO animals display a more modest shift (Figure 2B, C), suggesting an impairment in utilizing prior information - a feature of ASD and schizophrenia in humans^12 13 14^. Parameters of individual fits of the psychometric curves reveal specific changes in bias, with no changes in the other parameters (Figure 2D). To further examine their impaired utilization of priors, we measured how quickly mice adapt their choices following a block switch, when stimulus-reward contingencies change and updating of internal priors is required for optimal performance. We found KO animals are slower to update their priors following a block switch (Figure 2E). To quantify the latency of performance recovery, we fit a parametric curve to each animal’s performance trajectory, demonstrating significantly longer tau values (*p* = 0.004) and lower initial performance (*p* = 0.041) immediately after the switch in KO animals compared to WT littermates (Figure 2F, G).

**Figure 2.**
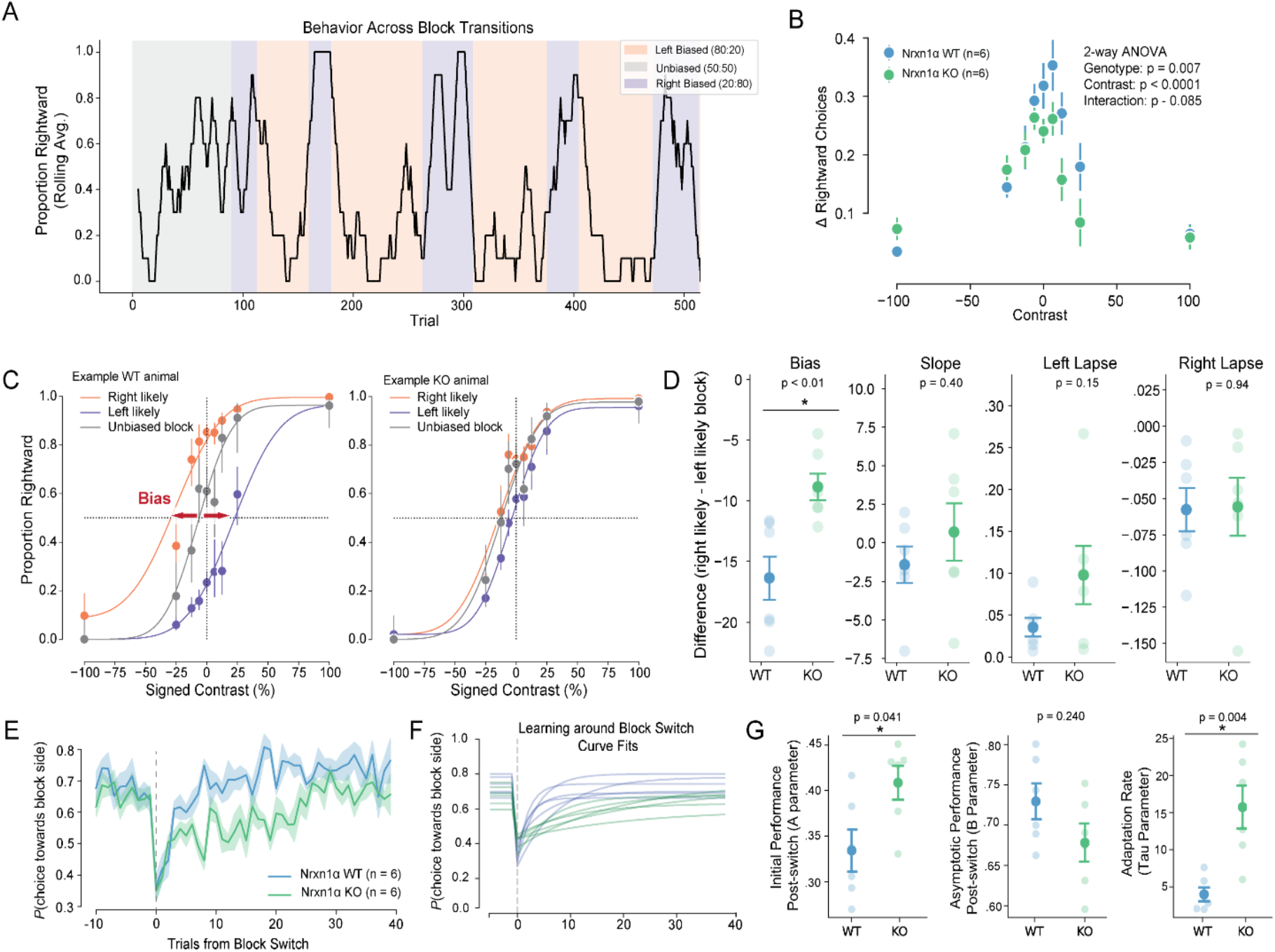
Nrxn1α mice exhibit slower adaptation to shifting reward contingencies. **(A)** Rolling average of a 10 trial window across a session depicting an animal’s choices conditioned on block probability. Shaded background indicates block-type, grey for 50:50 probability, coral for 80:20 stimulus probability, and purple for 20:80 stimulus probability. **(B)** Change in rightward choices between biased blocks, plotted by signed contrast. Nrxn1alpha KO animals showed reduced modulation across contrasts. **(C)** Psychometric functions from representative WT (left) and KO (right) animals across blocks with different priors. Curves are fit to data from left-biased (blue), right-biased (orange), and unbiased (gray) blocks. **(D)** Group-level comparisons of key psychometric parameters (bias, slope, lapse rates) across genotypes. Each point shows the within-session difference between left-biased and right-biased blocks (0.2 – 0.8). Error bars: ±1 SEM. **(E)** Mean proportion of probability of choices towards block side (e.g. right choices during right likely block and left choices for left likely block). WT mice adapt more strongly and consistently than KO mice. **(F)** Fits of a parametric model to performance around block transitions. Each line represents one session. **(G)** Summary of parametric fit parameters: initial accuracy (A), asymptotic performance (B), and adaptation rate (Tau). KO mice exhibit lower B and slower Tau. Error bars: ±1 SEM.

### Widefield calcium imaging reveals altered cortical dynamics in Nrxn1α KO mice during task performance

To investigate cortex-wide dynamics during decision-making, we crossed the Nrxn1α line onto the CamKII-tTA-tetO-GcaMP6s transgenic line ^15^, permitting widefield calcium imaging during behavior (Figure 3A). This approach provides the opportunity to survey cortex-wide neural dynamics in an unbiased manner ^16^. Imaging data was aligned to the Allen Mouse Common Coordinate Framework v3 (see Methods). Task-evoked responses in the visual cortex (VISp) showed robust activation following contralateral visual stimuli (Figure 3B). To examine specific task periods, we aligned the imaging to two events, stimulus onset and feedback (reward delivery or incorrect tone). This allows us to separately analyze the period of time pre-stimulus onset (inter-trial interval (ITI)), stimulus, and feedback periods. First, we examined baseline-corrected fluorescence (ΔF/F) for these task periods. During the ITI, increased activity was most prominent in motor regions in the Nrxn1α KO animals. In contrast, subsequent task periods including stimulus onset and feedback, neural activity spread broadly across cortex (Figure 3C). These maps suggest that Nrxn1α KO mice engage cortical networks more strongly and more diffusely than their WT littermates. Next, we asked whether this hyperactivity was accompanied by altered functional coupling between cortical regions. To test this, we computed pairwise correlations between regional ΔF/F time series, concatenated across trials. Nrxn1α KO mice showed globally increased correlations across cortex during the task indicating that cortical networks are tightly coupled (Figure 3D). Lastly, we assessed trial-to-trial variability, by computing pixel-wise variance maps across the three task periods. Nrxn1α KO mice exhibited markedly higher variance across all cortical regions and task periods compared to WT mice (Figure 3E). Elevated pairwise correlations in the Nrxn1α KO animals suggest greater global synchrony, while increased pixel-wise variance indicates larger moment-to-moment fluctuations in cortical activity. Altogether, these patterns are consistent with an increase in network gain, with the Nrxn1α KO cortex-wide neural activity being both globally synchronized and noisier.

**Figure 3.**
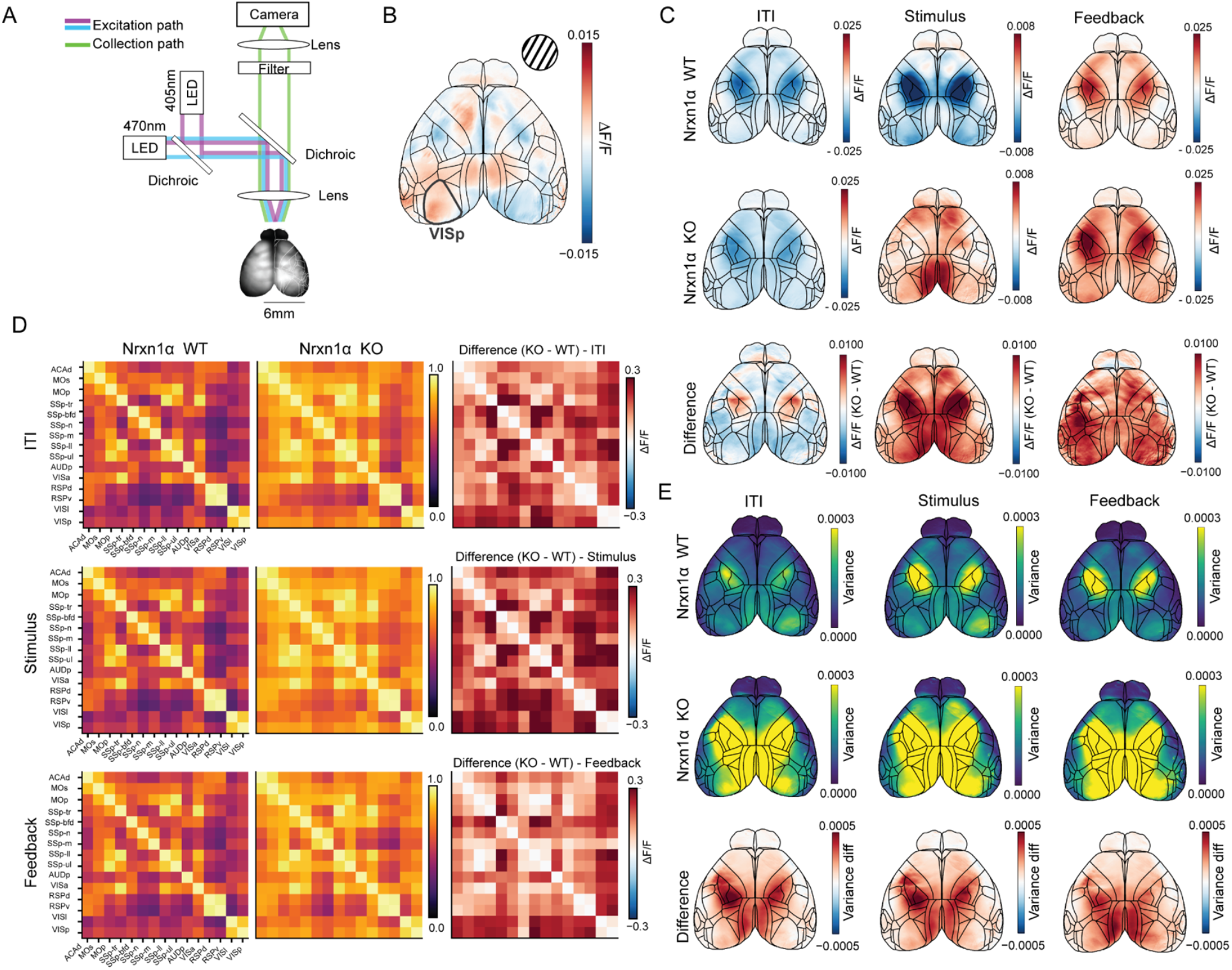
Widefield calcium imaging reveals altered cortical dynamics in Nrxn1α KO mice during task performance. **(A)** Schematic of the widefield imaging setup used to record cortical activity during behavior. Excitation light at 470 nm and 405 nm was delivered via LEDs, and emitted fluorescence was collected through a dichroic lens and detected by a camera. **(B)** Left: Average stimulus-evoked ΔF/F response map across all trials from a representative mouse. Gray outlined region overlay highlights primary visual cortex (VISp), which shows robust stimulus responses. **(C)** Trial-averaged ΔF/F maps across the cortex during three task epochs: Intertrial Interval (ITI), Stimulus, and Feedback. Top row: Nrxn1α WT mice. Middle row: Nrxn1α KO mice. Bottom row: KO–WT variance difference maps reveal regions of elevated variability in KO animals. **(E)** Pairwise correlations of regional activity (ΔF/F) during the task for WT (left) and KO (middle) mice. Right: Difference matrix (KO–WT) showing reduced inter-regional correlations in KO animals. **(E)** Trial-averaged variance maps across the cortex during three task epochs: Intertrial Interval (ITI), Stimulus, and Feedback. Top row: Nrxn1α WT mice. Middle row: Nrxn1α KO mice. Bottom row: KO–WT variance difference maps reveal regions of elevated variability in KO animals.

### An encoding model reveals unique contributions of movement variables to cortical activity in Nrxn1α KO mice

Previous work has demonstrated that widespread movement-related neural activity across cortex, including cognitive, sensory, and motor areas is largely dominated by movement signals ^17 18,19^. To understand how task and movement variables contribute to neural activity, we fit a linear encoding model incorporating task variables and multiple measures of movement (Figure 4A). Model performance, evaluated via cross-validated R^2^, was reassuringly high for both cohorts of mice, accounting for more than half of the total observed single trial variance. We took advantage of this strongly predictive model to separately evaluate the contribution of task variables and movements (Methods). Task variables included stimulus, choice, outcome and block probability. Movement variables included wheel velocity, and movements tracked by three cameras positioned to capture the left and right side of the mouse, motion energy of specific ROIs (e.g. whisker pad, base of tail), and a low dimensional summary (the top 200 dimensions from a singular value decomposition, SVD) of the three cameras positioned to capture the left/right side and body of the animal (see Supplementary Table 1 for a full list of variables).

**Figure 4.**
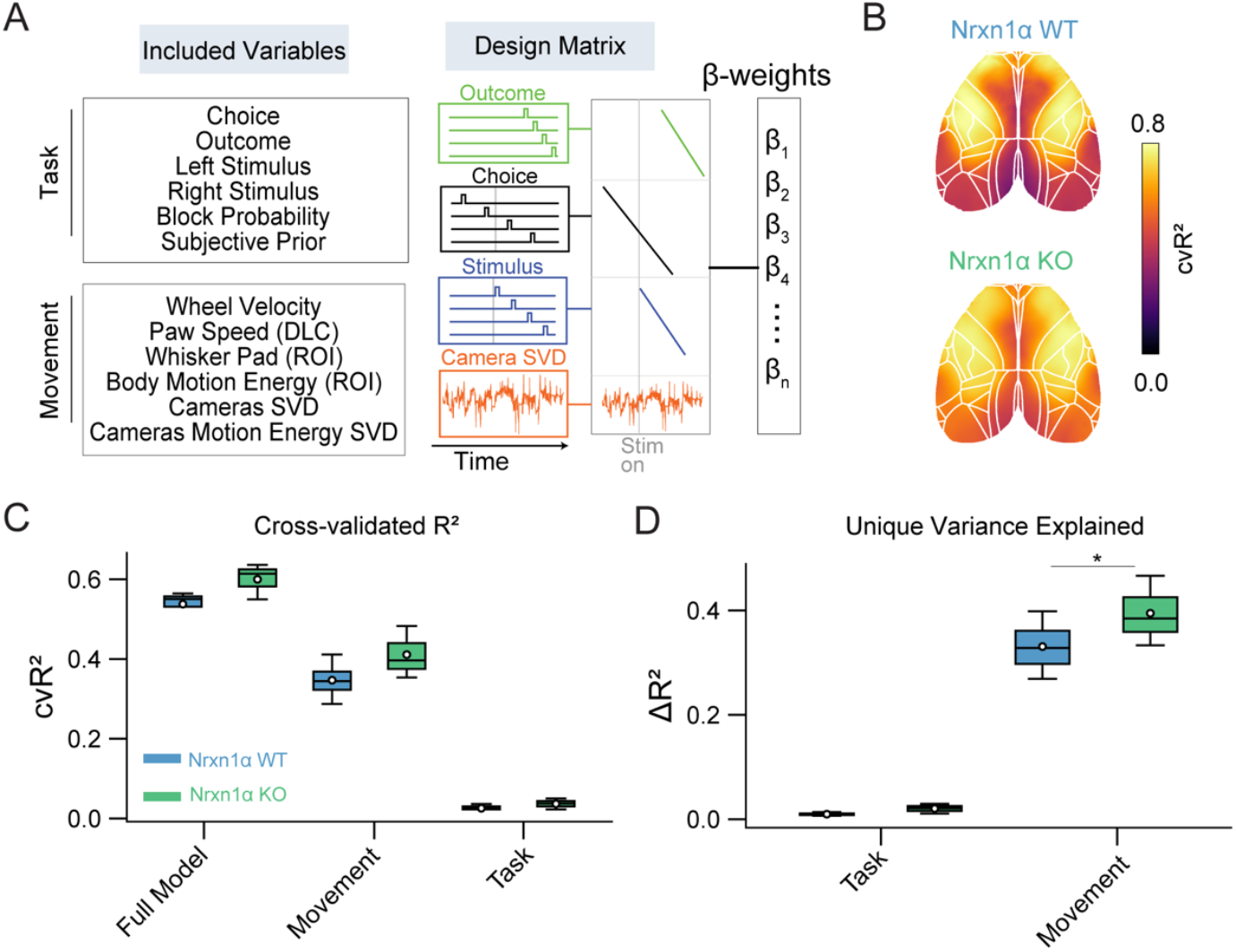
Linear encoding model reveals distinct contributions of movement variables to cortical activity in WT and Nrxn1α KO mice. **(A)** Schematic of the linear encoding model. The design matrix includes task-related variables (choice, outcome, stimuli, block probability, subjective prior) and movement-related variables (wheel velocity, paw speed from DeepLabCut, region-of-interest [ROI] motion energy from whisker pad and body, and a lower dimension representation, SVD, from multiple camera views). Each predictor is convolved with a temporal kernel, and β-weights are estimated to predict widefield neural activity. **(B)** Example maps of cross-validated R-squared (cvR^2^) averaged across animals, showing the proportion of cortical variance explained by the full model in WT (top) and Nrxn1α KO (bottom) mice. Outlines indicate Allen CCF region boundaries. **(C)** Group comparisons of cvR^2^for the full model, task regressors, and movement regressors, showing that movement-related predictors account for the majority of explained variance in both genotypes. Full model shown here is with trial time regressor subtracted out. WT (blue) and KO (green) distributions are plotted as boxplots with individual data points. **(D)** Unique contributions (ΔR^2^) of task and movement groups, quantified as the drop in explained variance when each group of regressors was shuffled.

We compared model performance across groups using both cvR^2^(variance explained by the full and shuffled models) and ΔR^2^(reduction in explained variance when predictors were removed) (Figure 4E, F). For cvR^2^, the full model explained substantial variance in both groups, with slightly higher cvR^2^in KO mice relative to WT (KO: 0.60 ± 0.02, WT: 0.54 ± 0.02; p = 0.076). Shuffling movement predictors produced a large reduction in CVR^2^(β = –0.40, p < 2 × 10−^16^), whereas shuffling task predictors had little effect (β = –0.02, p = 0.30). Importantly, there was a significant Group × Movement interaction (β = 0.071, p = 0.006), indicating that KO mice lost more explained variance when movement predictors were shuffled, consistent with greater movement dependence. No group differences were observed for task predictors (p = 0.70). ΔR^2^analysis confirmed this pattern: removing movement predictors reduced variance explained much more than removing task predictors overall (β = –0.38, p < 2 × 10−^16^). Moreover, KO mice showed significantly higher ΔR^2^than WT for movement predictors (KO: 0.40 ± 0.02, WT: 0.33 ± 0.02; p = 0.032), whereas no group difference was observed for task predictors (p = 0.78). Together, these analyses indicate that neural activity in KO mice is more strongly dominated by movement-related predictors than in WT, with little evidence for group differences in task-related variance.

Supplementary analyses of individual regressors were consistent with this pattern. Movement and camera-derived variables explained the largest share of variance (Supplementary Figure 3A, for movement variables, the green bars were reliably higher than the blue bars). To quantify the unique contribution of each predictor, we computed a unique explained variance (ΔR^2^,see Methods) for each variable (Methods). This is a more conservative estimate of each variable’s importance because it surmounts the problem of collinearity of regressors. We find Nrxn1α KO mice generally showing higher values (Supplementary Figure 3B). By contrast, task variables explained little unique variance in either group. Together, these results show that cortical activity in KO mice is disproportionately dominated by movement-related predictors, with minimal group differences in task-related encoding.

## Discussion

Impaired utilization of prior knowledge has emerged as a computational hallmark of neuropsychiatric disorders^9^.These atypical inference processes have been hypothesized to stem from disruptions in the underlying neural circuitry responsible for integrating past experience with current evidence. Our results show that Nrxn1α loss-of-function leads to impaired utilization of prior knowledge during a visuomotor task consistent with three other genetics models of ASD - Fmr1, Cntnap2, and Shank3B. Furthermore, we find that Nrxn1α KO mice show increased cortex-wide correlations in their neural activity. This heightened synchrony, coupled with increased variance explained by movement variables, points to a shift in cortical dynamics toward more global, possibly motor-dominated, activity patterns. Thus, this study reveals how disruption of synaptic signaling through Nrxn1α deletion impairs the integration of prior expectations into decision-making, leading to altered behavioral flexibility and widespread changes in cortical dynamics.

Recent work has demonstrated that mice flexibly incorporate learned priors into their decisions, and that this behavior can be captured using heuristic models of action history. The encoding of the subjective prior, the animal’s internal belief about the likelihood of stimulus side estimated by the model, was shown to be encoded in ∼20-30% of brain regions, spanning early sensory areas to motor regions ^20^. Furthermore, it was shown that a shift in weighting of prior encoding from sensory to frontal cortices mediates inflexible updating of priors in three genetic models of ASD (i.e. Fmr1, Cntnap2, and Shank3B) ^9^. These findings provide a fundamental basis for understanding how expectations shape behavior and how this process goes awry in neurodevelopmental disorders. Our study builds on this framework by examining how priors are represented at the neural population level across the cortex. By leveraging widefield calcium imaging, we were able to measure simultaneous activity across multiple cortical regions during decision-making, enabling us to assess how prior information is distributed and coordinated across brain areas. We find that Nrxn1α mutant mice show broadly elevated cortical activity and increased variance explained by movement, suggesting that disruptions in cortex-wide neural activity and movement-related signals could interfere with the neural mechanisms supporting prior-dependent decision-making. These results extend Noel’s findings by showing that Nrxn1α loss alters large-scale cortical activity and movement-related dynamics, revealing a novel mechanism through which this gene can impact neural circuit function.

Although overall model performance (cvR^2^) was high for both WT and KO mice, indicating that the encoding model captured neural activity well for both groups. However, our analysis of the variance uniquely explained by each variable (ΔR^2^) revealed genotypic differences. Nrxn1α KO mice showed a significantly larger reduction in explained variance when movement regressors were shuffled, suggesting that their cortical activity was more tightly coupled to movement variables. While an overall increase in movement may in part explain these results (Supplementary Figure 2), the encoding model accounts for movement magnitude, therefore the larger ΔR^2^ effect likely reflects a qualitative shift in how movements are represented across cortical populations. There are a few explanations for these results, (1) stronger coupling between sensory and motor areas, (2) an increase in stereotyped movements resulting in predictable movement-related neural activity, or (3) hyperconnected cortical circuits may amplify the impact of movement-related signals leading to stronger dominance of motor activity. Overall, these findings suggest a reweighting of cortical computations in the Nrxn1α KO animals, where widespread hyperconnectivity may bias the network toward motor-driven dynamics at the expense of task-related processing.

States of engagement have previously been shown to impact neural activity underlying decision-making^21^. Mice in disengaged states displayed higher trial-to-trial neural variability and stronger encoding of task-independent movements, even though their overall movement magnitude remained unchanged. In our case, increased explained variance by movement variables in Nrxn1α KO mice could reflect altered engagement dynamics - a shift toward more variable, less task-coupled behavior - rather than purely movement driven cortical states. Furthermore, differences in frequency in transitions between engagement states could lead to deficits in maintaining stable states necessary for integrating prior information effectively. Although movement-related signals explain more variance in Nrxn1α KO mice, we cannot determine whether this reflects a causal shift in behavioral strategy or a consequence of underlying neural disorganization. Disentangling whether increased movement-related activity reflects compensatory strategies, reduced task engagement, or fundamental alterations in circuit processing would require manipulating cortical state or movement structure to establish directionality.

In summary, we show Nrxn1α loss-of-function leads to impaired utilization of prior knowledge when there are changes in the environment. The increased variance explained by movement variables and elevated inter-regional correlations in Nrxn1α KO mice point to a shift toward more globally synchronized, movement dominated cortical states. Together, these results provide insight into how neural dynamics are shaped across multiple regions when combining sensory evidence with prior knowledge and how this process is modulated by a disease-associated perturbation.

## Methods

All experiments were conducted in accordance with the National Institute of Health Guidelines for the Use of Animals, and all procedures approved by the Institutional Animal Care and Use Committee of the University of California at Los Angeles. Animals used in experiments had not previously been involved in other experiments or exposed to any drugs. Mice were housed under a 12-h light/12-h dark reverse light cycle and provided food *ad libitum*. All mice are housed with enrichment in the form of mouse igloos with fast-tracs (Bio-Serv) and yogurt drops or peanuts.

### Animals

Constitutive Nrxn1α KO mice were obtained from the Fuccillo lab (University of Pennsylvania) and were kept on a C57BL/6 background (Geppert et al. 1998). To transgenically express GcaMP6s, Nrxn1α +/-animals were subsequently crossed onto TRE-GCaMP6s (G6s2, JAX 024742) (Wekselblatt et al. 2016) and mice expressing tTA under the control of the alpha isoform of calcium/calmodulin-dependent protein kinase type II alpha chain (CaMKII) promoter (JAX 003010). Offspring were weaned at P21 and separated by sex in cages of 2-5 animals of mixed genotypes. Wild-type littermates were used as controls. Adult male and female mice between the ages of 2-6 months were used for experiments.

### Behavioral Training

Mice were habituated and trained as described previously ^10,20^.

#### Habituation

Once mice begin water-restriction, they are handled for 5-10 minutes for three days to acclimate mice to human interaction. To reduce stress, mice are gently scooped by one or two hands rather than being lifted by the tail. During handling, mice are given 1mL water from a syringe while restraining their headbar for short periods of time (10 seconds). On the fourth day, mice explore the behavioral rig and are given water from the rig’s reward spout. After this the mouse is head-fixed for three days (1st day: 20 minutes, 2nd day: 40 minutes, 3rd day: 60 minutes) and is given auto-rewarded trials. In these trials, the wheel is immovable (secured with tape) and the Gabor patch (100% contrast, 0.1 cycles/degree spatial frequency, random phase, vertical orientation) is presented at one of two random initial locations, -35° (left), +35°(right) for an average of 10 seconds. The Gabor patch then appears in the center of the screen, and a 3μl water reward with 10% sucrose is given after 500ms. If mice receive less than 1mL of water or their weight drops below 85% of baseline, they are supplemented at the end of the day with water or diet gel.

#### Training

Following the habituation protocol, the wheel is unlocked and coupled to the movement of the visual stimulus. The visual stimulus appears on either the left or right side of the screen (+/-35° azimuth, within the field of binocular vision in mice^22^). For a correct response, the mouse must move the wheel to center the visual stimulus, while moving the stimulus off the screen is considered an incorrect response. If the mouse incorrectly responds or no response is recorded for 60 seconds, then the mouse receives a time-out where the Gabor patch is fixed at 70° azimuth for 2 seconds on the side the stimulus originated on, and white noise is played for 500ms. To begin a trial, the wheel must not move for a quiescence period of 400-700ms (randomly drawn from an exponential distribution with a mean of 550ms). If the wheel was moved during this period, the quiescence period is restarted. As mice are learning the basic task, the stimulus appears equally on either side (50/50). There are 6 stages of training described below:

Phase 1: Only 50% and 100% contrasts are presented. To progress to the next stage, mice must perform at or above 80% correct for each contrast on both sides calculated over the previous 50 trials. Phase 2: 25% contrast is added. To progress, same criterion as phase 1.

Phase 3: 12.5% contrast is added. To progress, the mouse must complete 200 trials irrespective of performance.

Phase 4: The 6% contrast is added. To progress, the mouse must complete 200 trials irrespective of performance.

Phase 5: 0% contrast is added, rewarded equally on either side in a random manner. To progress, the mouse must complete 200 trials irrespective of performance.

Phase 6: 50% contrast is dropped from the set, and repeat trials are dropped. Final contrast set: 100, 25, 12.5, 6.25, 0.

After mice reach proficiency on the 50/50 version of the task (criterion described in^10^), they are moved to a block structure task. The probability of the Gabor patch is presented on the left or right with a probability determined by the block, 80:20, left likely block or 20:80, right likely block. Block length ranges from 20-100 trials.

### Surgical Procedures

All surgeries were performed on a stereotaxic frame under isoflurane anesthesia (3% for induction; 1-2% during surgery). After induction of anesthesia, 1.2 mg/kg of meloxicam was injected subcutaneously and eyes covered with ophthalmic ointment. Sterile surgical technique was used throughout. Fur over the skull was removed using a small electric hair trimmer, followed by a depilatory cream. The surgical area was then cleaned with alternating applications (3 times) of 70% isopropyl alcohol and betadine. Four small incisions were made in the scalp to expose the skull. The periosteum is gently removed using a scalpel blade for proper optical clearance. Vetbond is then applied to the exposed skin 1mm around the cuts. Using a small amount of cyanoacrylate glue (Zap-A-Gap CA+, Pacer technology), the headplate was fixed to the skull. Dental cement (C&B Metabond, Parkell) was then applied to further secure the headplate. A 3D printed black cone was placed over the headplate and surrounded the exposed skull. Open areas where the cone met the tissue were filled with dental cement (Ortho-Jet, Lang Dental). Subsequently, a thin layer of cyanoacrylate glue is applied, repeating three times after each layer is cured. After all four layers of cyanoacrylate glue have cured, cortical blood vessels are clearly visible^16^. Animals are monitored for 72 hours and are given meloxicam (1.2mg/kg) and enrofloxacin (5 mg/kg) subcutaneously.

### Widefield Imaging

Widefield imaging was carried out using an inverted tandem-lens macroscope, paired with an sCMOS camera (Edge 5.5, PCO) operating at 60 frames per second. In this setup, the objective lens is inverted—meaning the camera mount faces the brain surface—to project near-collimated light onto a second lens, which then focuses the light onto the camera sensor. The optical system included a 105 mm focal length top lens (DC-Nikkor, Nikon) and an 85 mm bottom lens (85M-S, Rokinon), yielding a total magnification of 1.24x. This configuration provided a field of view measuring 12.5 × 10.5 mm, with images acquired at a resolution of 640 × 540 pixels after applying 4x spatial binning, resulting in an approximate spatial resolution of 20 μm per pixel.

To detect GCaMP fluorescence, a dichroic mirror was used to direct blue (470 nm) and violet (405 nm) excitation light onto the sample while allowing the emitted longer-wavelength light (>495 nm) to pass through a green bandpass filter (525 ± 25 nm, #86-963, Edmund Optics) placed in front of the camera. Excitation light was projected onto the cortical surface using a 495 nm long-pass dichroic mirror (T495lpxr, Chroma) positioned between the two macro lenses. A collimated blue LED (470 nm, M470L3, Thorlabs) and a collimated violet LED (405 nm, M405L3, Thorlabs) were coupled into the same optical path with a dichroic mirror (#87-063, Edmund Optics).

Illumination alternated between the two LEDs on a frame-by-frame basis, producing a sequence of frames excited by blue light and another by violet light, each recorded at 30 fps. The blue-excited frames captured calcium-dependent fluorescence, while the violet-excited frames recorded fluorescence independent of calcium activity. By capturing both signals, we enabled offline hemodynamic correction to account for intrinsic signals such as hemodynamic responses. For this correction, the violet-excited fluorescence was rescaled to match the intensity of the blue-excited fluorescence and then subtracted to remove non-neural components, thereby isolating the true calcium-dependent signal. All subsequent analyses were based on this processed differential signal at 30 fps.

During imaging, three additional cameras captured movement: Camera 1 (Left): 60Hz, full resolution (1280×1024), Camera 2 (Right): 150Hz, half resolution (640×512), and Camera 3 (Body): 30Hz, half resolution (640×512).

### Preprocessing of Widefield Calcium Imaging

Widefield imaging data were preprocessed using the wfield pipeline (https://github.com/jcouto/wfield). Manual alignment to the Allen reference atlas was performed using four anatomical landmarks. For motion artifacts, raw binary data were first motion-corrected by 2D rigid registration to the mean of all frames, aligning frames to correct for drift. Subsequently, to calculate ΔF/F, a baseline image was computed by averaging a set number of initial frames, and this was subtracted from each frame to normalize fluorescence activity. The resulting data were decomposed via singular value decomposition (SVD), retaining the top 200 components for dimensionality reduction. This separates the data into temporal and spatial components. Hemodynamic artifacts were removed using a pixelwise regression to isolate and correct for blood flow-related signals. This was done by first filtering with a 0.1 Hz temporal high pass filter, scaling the 405nm signal to the 470nm signal, and then subtracting it out.

### Linear Encoding Model

To relate behavioral variables to neural activity, we employed linear encoding models previously used in our lab^17,23^. We constructed a design matrix that included a range of task and movement-related variables, allowing us to quantify how each contributed to neural dynamics. The model was designed to capture both fixed and time-varying relationships between behavior and neural activity across the trial timeline.

The design matrix incorporated discrete behavioral events (e.g., stimulus onset, choice, and reward delivery) using time-aligned binary regressors that were set to 1 at the time of the event and 0 otherwise. To capture the temporal influence of events like choice and outcome, we included time-shifted versions of these regressors that spanned the trial duration. A time-varying intercept regressor was also included—this consisted of a set of binary vectors with a 1 at the same time point across all trials, accounting for baseline fluctuations in activity across the trial.

We included the following variables as regressors:

– Task variable regressors: choice, right/left stimulus, outcome, true prior (determined by the task’s block structure), and subjective prior (estimated using the ActionKernel model^10^).
– Instructed movement regressors: wheel velocity (measured via a rotary encoder), paw speed (measured using DeepLabCut).
– Uninstructed movement regressors: lower dimensional representations of the full behavioral video (Video SVD) and the motion energy (motion energy SVD) of all three cameras (left, right, and body). We also included motion energy from predefined regions of interest (e.g., whisker pads and body) using pixel-based coordinates.

To prevent overfitting, we implemented ridge regression with L2 regularization. Model weights were trained on 90% of the data, with the remaining 10% reserved for testing. To assess overall model performance, we performed 10-fold cross-validation to compute explained variance (cvR^2^). Additionally, we quantified the unique contributions of individual variables using a reduced model approach. For each variable, we fit a model in which only that variable’s regressors and the time-varying intercept were left intact, while all other regressors were temporally shuffled. This ensured that only the unshuffled variable could account for structured variance in the neural activity. We then also fit an intercept-only model. The unique explained variance (ΔcvR^2^) for each variable was then computed as the difference in cross-validated R^2^between the single-variable model and the intercept-only model. This allowed us to isolate the variance explained by each behavioral or movement variable while controlling for trial-aligned temporal structure. To further assess the contributions of different variable groups, we performed grouped shuffles of the linear encoding model. Specifically, we created one model in which all task-related regressors were temporally shuffled, and another model in which all movement-related regressors were shuffled. For each shuffle, we calculated the change in cross-validated R^2^relative to the full model, providing a measure of the variance uniquely attributable to the shuffled variable group. This approach allowed us to evaluate the impact of task versus movement variables on neural activity.

### Data Reporting and Statistical Analyses

Unless otherwise indicated, data are presented as mean ± standard error of the mean (SEM). Statistical comparisons were performed using two-way analysis of variance (ANOVA) followed by appropriate post hoc tests when indicated in the figure legends. Linear mixed-effects models were fit in R (Imertest) to account for repeated measures across animals and sessions. All other statistical analyses were performed in Python (SciPy, statsmodel). The threshold for statistical significance was set at *p* < 0.05. Exact statistical tests, sample sizes (n), and *p* - values are reported in the figure legends.

## Data/code availability

All the code used for analyses will be made public upon publication. Packages used in this project include: LabCams for camera acquisition (https://github.com/jcouto/labcams), wfield for preprocessing and analytic tools (https://github.com/jcouto/wfield) and linear encoding analyses (https://github.com/churchlandlab/chiCa).

## Acknowledgements

We thank Anup Khanal for his guidance in setting up the behavioral rig and support with IBL infrastructure. We also thank Mayo Faulkner for assistance with setting up the pipelines for handling the widefield calcium imaging data within the IBL. We also acknowledge initial support within the IBL from Miles Wells, Gaelle Chapuis and Olivier Winter. We are grateful to all members of Churchland lab for their valuable discussions and for sharing common resources. This work was supported by NIH grant U19NS123716 (to A.K.C), including a diversity supplement that supported M.F.D. M.F.D was also supported by a Burroughs Wellcome Fund Postdoctoral Enrichment Program (PDEP) award.

## Author contributions

M.F.D. and A.K.C. conceived and designed the project. M.F.D performed all surgeries, collected the imaging and behavioral dataset, and carried out data analysis. J.C. built the widefield imaging, developed analysis tools, and preprocessed the imaging data. M.M. assisted in constructing the behavioral rigs, provided analytical guidance. L.T.O. ported the linear encoding model to Python and provided guidance in model implementation. C.F. modified the behavioral models. P.M. and I.H. assisted with behavioral training of the mice. A.K.C supervised the project and secured funding. M.F.D and A.K.C wrote the manuscript.

---

**Supplementary Figure 1.**
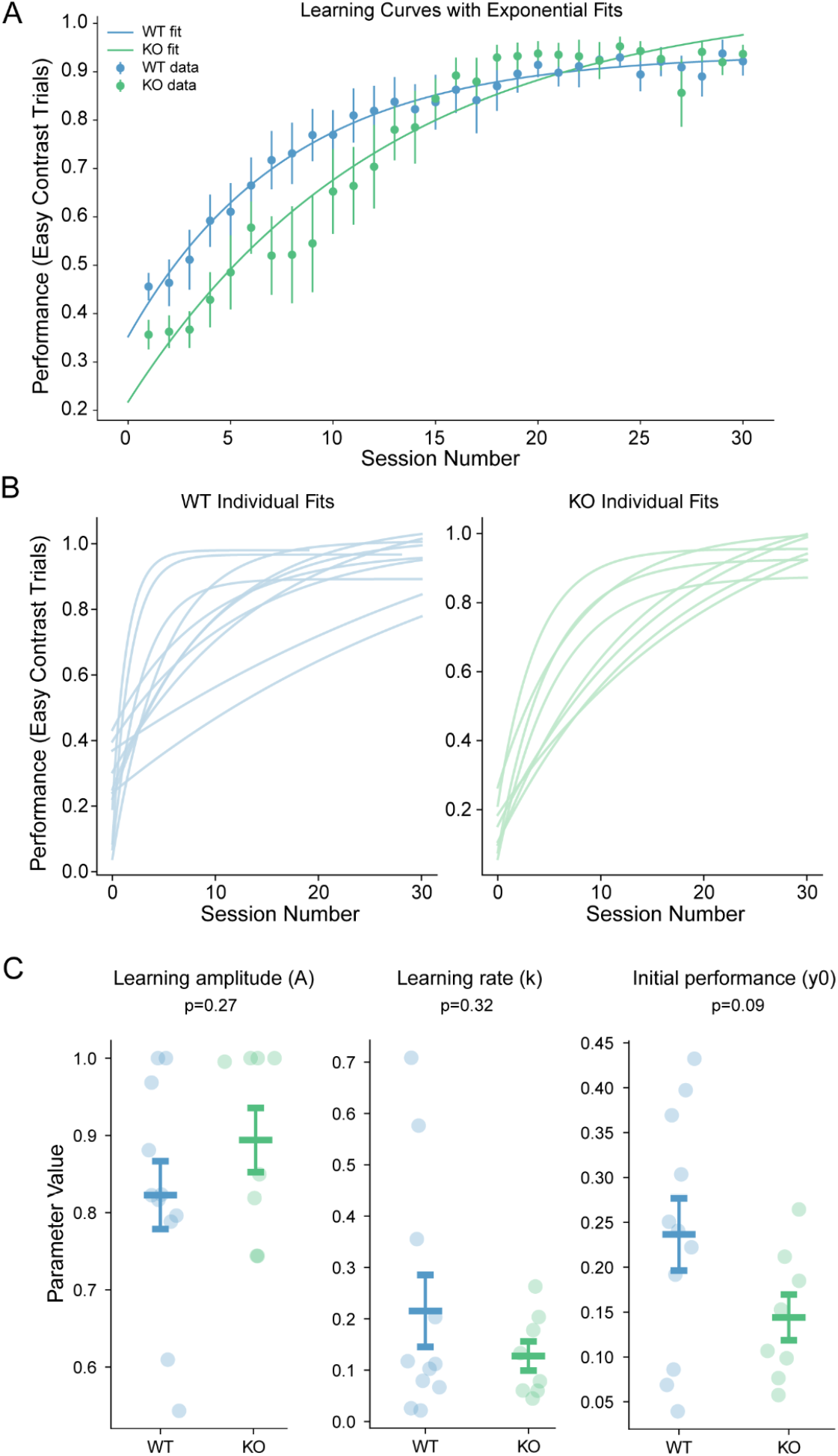
Learning curves and exponential fits are comparable between Nrxn1α WT and KO mice. (A) Group-averaged performance on easy contrast trials (contrast set: 50, 100) across training sessions. Dots represent mean ± SEM for WT (blue) and KO (green) mice. Sold lines show exponential fits for the data. (B) Exponential fits to individual animals’ performance curves. Left: WT mice; Right: KO mice. Each line represents one animal. (C) Parameters extracted from exponential fits. Left: learning amplitude (A), middle: learning rate (k), right: initial performance (y0). Each point corresponds to one animal, with mean ± SEM shown. P - values indicated group comparisons (WT vs KO).

**Supplementary Figure 2.**
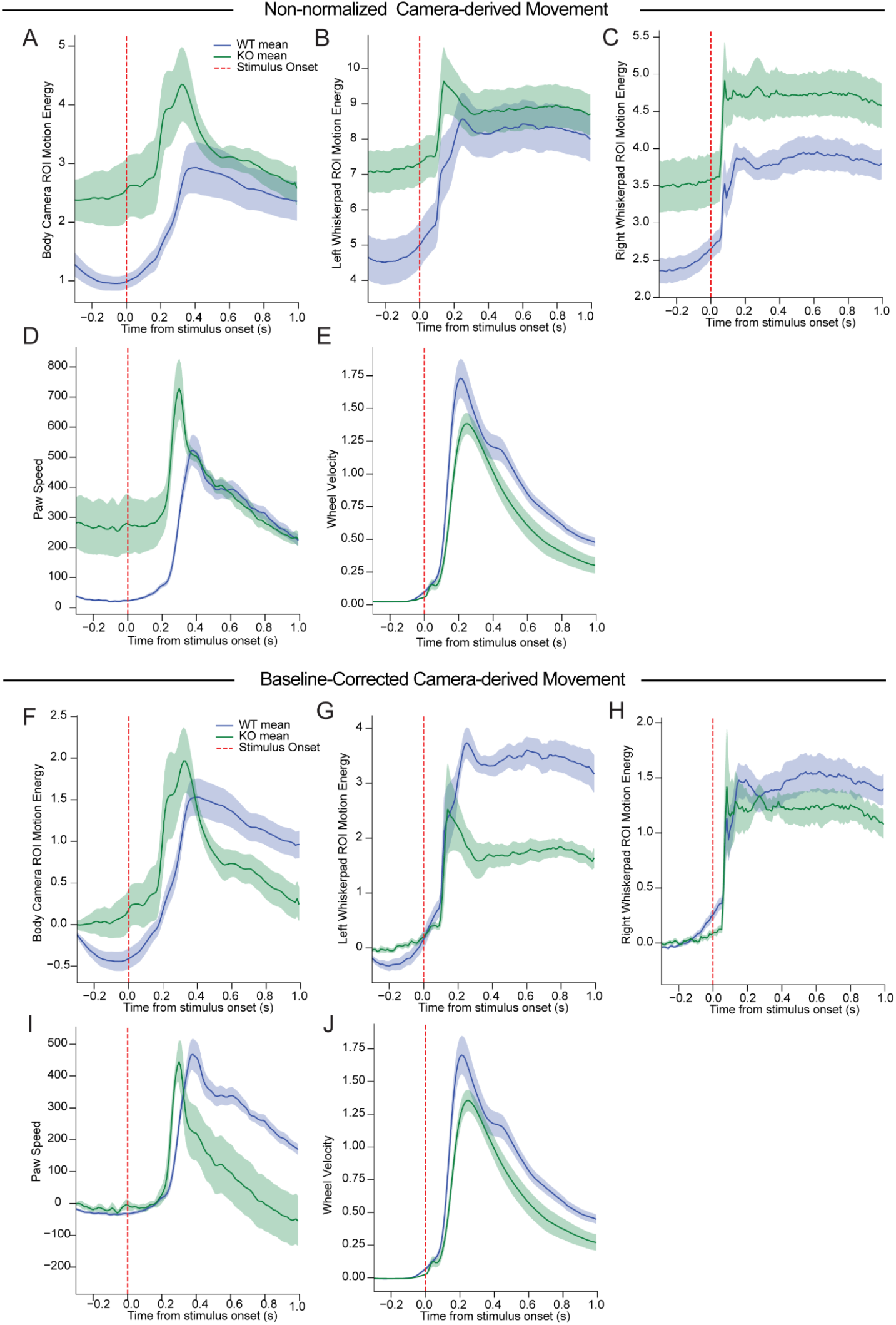
Camera-derived movement measurements in Nrxn1α WT and KO mice. (A-E) Non-normalized movement traces aligned to stimulus onset (red dashed line). (A-C) Body and whisker pad motion energy quantified from camera-derived regions of interest (ROIs). KO mice exhibited elevated baseline motion energy relative to WT. (D) Paw speed was extracted from DeepLabCut (DLC) tracking of paw position, and (E) wheel velocity was measured using a rotary encoder. (F-J) Same as (A-E), but with baseline-corrected signals (subtraction of pre-stimulus mean). Baseline correction reveals comparable stimulus-evoked dynamics between genotypes across all measures. Shaded areas represent ± SEM.

**Supplementary Figure 3.**
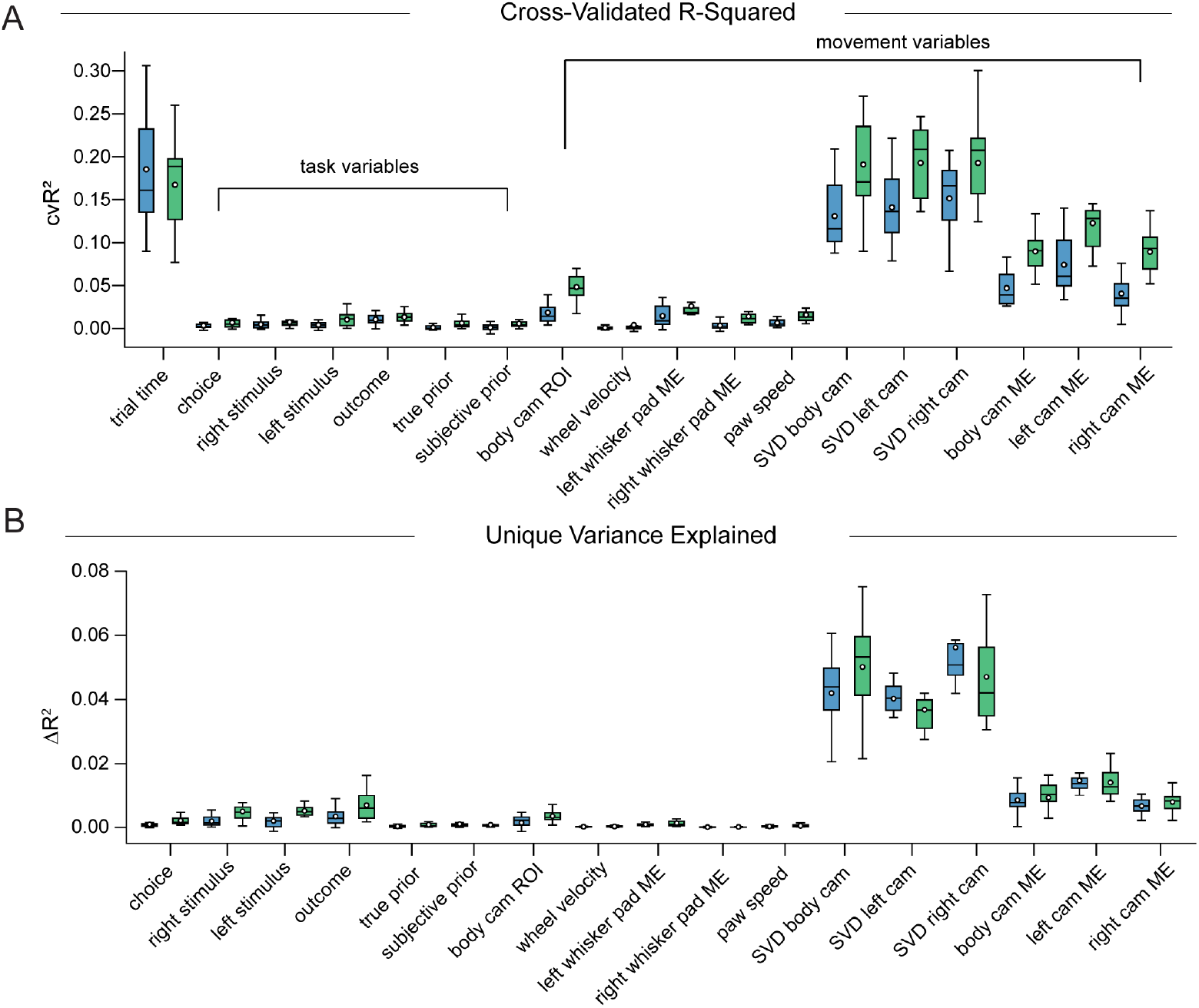
Cross-validated variance explained by task and movement variables. **(A)** Cross-validated R-squared (cvR^2^) values for each individual variable included in the linear encoding model, indicating how well each predictor explains cortical activity on its own. Task variables (choice, outcome, left/right stimulus, block structure, and priors) are shown on the left, while movement variables (wheel velocity, paw speed, whisker pad motion energy, body motion energy, and camera-derived components) are shown on the right. Each boxplot represents the distribution across animals (blue = Nrxn1α WT, green = Nrxn1α KO), with circles showing individual sessions. Movement-related regressors account for substantially higher variance compared to task-related regressors. **(B)** Unique contribution (ΔR^2^) of each variable, calculated as the drop in cross-validated R^2^when that variable is shuffled while all others remain intact. This approach isolates the variance uniquely explained by each predictor, controlling for shared temporal structure. Movement regressors show greater unique contributions to cortical activity compared to task regressors, highlighting the dominant influence of spontaneous and uninstructed movement on widefield signals.

**Supplementary Table 1.**

| Variable name | Description | Variable Type | Category |
| --- | --- | --- | --- |
| Choice<br>(Left/Right/NoGo) | All frames in either a left - or<br>a rightward choice trial | Event Kernel | Task |
| Left Stimulus | All frames after a leftward<br>stimulus | Event Kernel | Task |
| Right Stimulus | All frames after a rightward<br>stimulus | Event Kernel | Task |
| Outcome | All frames after a successful<br>choice, rewarded | Event Kernel | Task |
| Block Probability (True<br>Prior) | Probability of stimulus side<br>given current block structure<br>(80/20, 20/80) | Event Kernel | Task |
| Subjective Prior | Animal's inferred prior<br>(estimated from behavior,<br>modeled by ActionKernel) | Event Kernel | Task |
| Body Cam ROI | Motion energy from body<br>camera ROI | Analog | Movement |
| Wheel Velocity | Wheel velocity (cm/s)<br>extracted from the rotary<br>encoder | Analog | Movement |
| Whisker pad Motion<br>Energy (Left/Right) | Motion energy from whisker<br>pad regions | Analog | Movement |
| Paw Speed | Speed of forepaw<br>movements (tracked via<br>DLC) | Analog | Movement |
| Camera Video SVD | Video dimensions from all<br>three cameras (body, left,<br>right), SVD | Analog | Movement |
| Camera Video Motion<br>Energy | Video dimensions from<br>motion energy for all three<br>cameras (body, left, right) | Analog | Movement |

